# Variant-resolved prediction of context-specific isoform variation with a graph-based attention model

**DOI:** 10.1101/2025.09.18.677148

**Authors:** Aviya Litman, Zhicheng Pan, Ksenia Sokolova, Joyce Fang, Tess Marvin, Natalie Sauerwald, Christopher Y. Park, Chandra L. Theesfeld, Olga G. Troyanskaya

## Abstract

In eukaryotes, most genes produce multiple transcript isoforms that diversify the transcriptome and proteome, serving as a key mechanism of functional regulation. Genetic variation can disrupt the RNA processing signals that shape isoform structure and abundance, yet modeling these effects at full-length isoform resolution remains challenging due to the complexity of transcript regulation. Here, we introduce Otari, an attention-based graph neural network framework trained on the human genomic sequence and long-read transcriptomes across 30 tissue types and brain regions. Otari predicts tissue-specific differential isoform abundance by integrating sequence-derived epigenetic and post-transcriptional signals, enabling isoform-resolved variant effect interpretation. Applied to large-scale variant datasets including an autism cohort, Otari uncovers patterns of isoform dysregulation undetectable at the gene level, such as variant-driven perturbations in isoform abundance and microexon usage implicated in autism pathophysiology. We provide Otari as a resource for powering isoform-level analyses across tissues at scale.

## Main text

Understanding transcript isoform diversity is essential for capturing the precise spatiotemporal expression of the genome in human health and disease^1–4^. This diversity arises from tightly regulated programs of isoform usage and switching across tissues and cell types, and their disruption has been implicated in a wide range of conditions^5–16^. Evaluating genetic variant effects at the isoform level can therefore uncover clinically relevant mechanisms and inform isoform-targeted therapies^17–20^.

Recent studies have shown that isoform-level analyses often reveal stronger effect sizes and greater disease specificity than gene-level analyses^8–10,21,22^. For example, transcriptome profiling of the cerebral cortex in autism, schizophrenia, and bipolar disorder identified differentially expressed transcripts not detected at the gene level, uncovering new candidate disease genes^10^. An integrative framework is needed to model full-length, context-specific isoform variation that arises from the complex interplay of splicing factors, regulatory elements, epigenetic modifications, and the spliceosome^2,3,12,23,24^. To this end, we developed Otari: a comprehensive and interpretable graph-based framework of isoform regulation, powering the characterization of transcriptomic diversity and isoform-level variant effects at scale.

While sequence-based deep learning models have been successful in a variety of biological and disease contexts^25–29^, extending these methods to full-length transcript isoforms remains a major challenge. Existing approaches in this space have made significant progress in predicting individual regulatory features, such as splice sites (e.g. SpliceAI, MMSplice, Pangolin)^30–35^ or RNA binding protein profiles (e.g. Seqweaver, RBPNet)^36,37^. However, they fall short in capturing the regulatory complexity of full-length isoforms and are unable to predict isoform abundance levels. These limitations can be attributed in part to the challenges of this modeling task, but also to constraints of short-read RNA sequencing, which lacks the resolution to span full transcripts or multiple splice junctions.

Our integrative graph-based framework, Otari, addresses the challenges of modeling full-length transcript isoforms at scale. Otari is trained entirely on long-read transcriptomic data, which provides direct measurements of full isoforms even in data-sparse settings^38^, and leverages advances in attention-based graph deep learning to capture whole transcript regulation. Transcript isoforms are naturally phrased as graphs, and are therefore well-suited for a message-passing graph neural network (MPNN) approach. Unlike convolutional or transformer-based models, MPNNs learn directly from dynamically structured graphs^39^ while leveraging attention mechanisms to capture both local and global regulatory signals shaping transcriptomic variation.

Otari effectively learns tissue-specific regulatory patterns across 30 human tissues and brain regions by embedding epigenetic and post-transcriptional regulatory signals into DNA sequence graphs, enabling accurate and explainable predictions of transcriptome-wide isoform-level abundance. We show that Otari generalizes to novel isoforms, complex multi-exonic transcripts, and across both protein-coding genes and long non-coding RNAs. We further extend the framework to predict variant effects on isoform abundance, enabling isoform-resolved variant interpretation at scale. Applied to thousands of variants from HGMD^40^, ClinVar^41^, and GTEx^42^, Otari reveals patterns of transcript-level variant effects that are undetectable at the gene level, providing hypotheses for previously unexplored regulatory mechanisms underlying human health and disease. Finally, in a case study of a large autism cohort^43^, Otari identifies variant-driven isoform misregulation, including altered microexon usage, as a prevalent feature of autism pathophysiology.

## Results

### Developing a graph-based model of isoform regulation

Otari is an attention-based graph deep learning framework that embeds full transcripts into structure-aware representations, enabling isoform-level predictions across tissues (Fig. 1a). Each transcript is represented as a directed graph, where nodes encode DNA sequence features for splice site regions and edges define exon connectivity, effectively modeling isoform splice structure. This graph-based design enables a range of downstream tasks, including prediction of isoform-resolved variant effects and context-specific analysis of isoform regulation (Fig. 1b).

**Figure 1.**
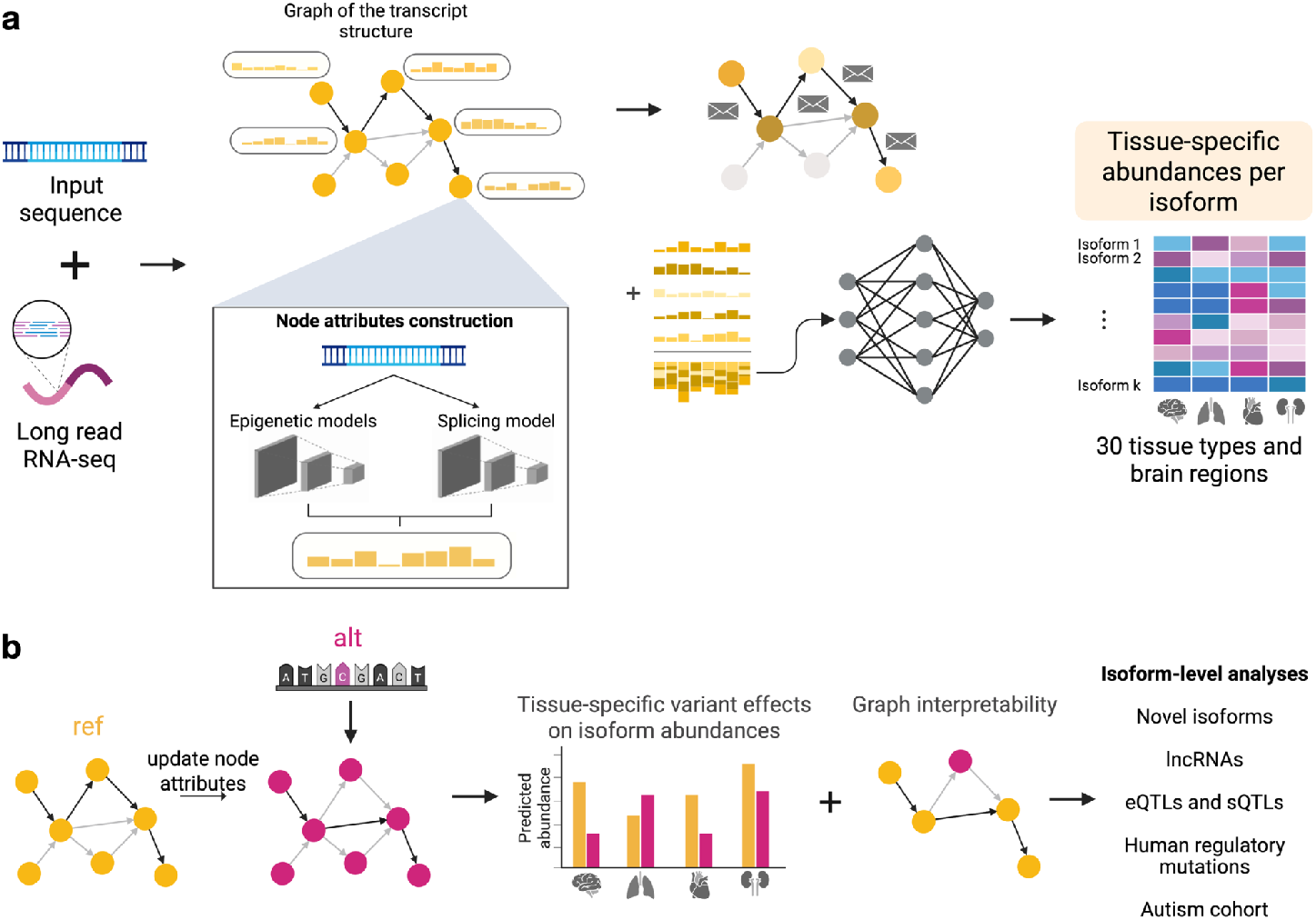
Overview of the Otari framework. **a**, Data construction and model training protocol. Transcript graphs were constructed from the human genomic sequence and long-read RNA-seq data. Nodes represent sequence features surrounding exon splice site regions, and directed edges encode exon connectivity based on GENCODE v47 annotations. Each node is assigned an attribute vector comprising outputs from three sequence-based models (ConvSplice, Sei, and Seqweaver) that capture 5′ and 3′ splice site region regulatory features. A graph attention network (GAT) propagates information across nodes through attention-based message passing. The resulting node embeddings are pooled into a graph-level representation and regressed onto a 30-dimensional vector of tissue-specific isoform abundances derived from long-read RNA-seq data. At inference, Otari outputs relative isoform abundances across tissues. **b**, Otari enables prediction of isoform-level variant effects. For a given variant, transcript graph node sequences are mutated and node attributes are updated. Variant impact is estimated as the log fold change between predictions from reference and alternative graphs. These predictions are combined with interpretability metrics to identify transcript nodes affected by *cis*-regulatory variation and impacted underlying regulatory features. Otari facilitates downstream analyses of isoform regulation across tissues and contexts.

Otari integrates the regulatory landscape of each isoform, captured by constructing rich node embeddings that include splicing, chromatin, and RNA-binding protein (RBP) profiles. To model splicing signals, we developed ConvSplice, a convolutional neural network for splice site strength prediction (Supp. Fig. 1a). ConvSplice achieved state-of-the-art performance, with precision-recall AUCs of 0.98 for both donor and acceptor site prediction, surpassing SpliceAI’s^30^ 0.97 (Supp. Fig. 1b). In top-k accuracy evaluation, ConvSplice achieved 0.94 for both site types, outperforming SpliceAI’s 0.93 and 0.92 with particularly improved accuracy at donor sites. The improved performance can be attributed to two factors: (1) the deeper architecture with 40 convolutional layers and (2) the larger 20kb sequence context that allows the model to capture longer-range sequence dependencies affecting splicing regulation. To incorporate epigenetic signals in our node embeddings, we used the Sei model^26^, trained on chromatin immunoprecipitation sequencing (ChIP-seq) data to predict tissue- and cell-type-specific *cis*-regulatory chromatin features. For post transcriptional regulation-based node embeddings, we leveraged Seqweaver^36^ to infer binding affinities of over 100 RBPs that modulate splicing, processing, transport, stability, and translation^44^.

These regulatory signals were embedded in a custom message-passing graph neural network (MPNN) designed to capture both local and global transcript features. Our architecture combines attention mechanisms, residual connections, and pooling layers (Supp. Fig. 2), and a systematic architecture and hyperparameter search confirmed that this design outperforms alternative models (Supp. Fig. 3).

### Otari predicts tissue-specific isoform abundance

We trained Otari to regress pooled transcript graph embeddings onto relative isoform abundances (measured in TPM) across 30 human tissues and brain regions^45^, and evaluated its performance in distinguishing high-versus low-expressed transcripts (Methods). Otari achieved strong predictive accuracy, with an average area under the receiver operating curve (AUROC) of 0.835 and precision-recall curve (AUPRC) of 0.812 on the chromosome 8 holdout test set from the Gao et al.^45^ dataset (Fig. 2a; mean Pearson’s *r* = 0.563; max *p* = 2.1 × 10^−94^), indicating that the model successfully captures the regulatory code underlying tissue-specific isoform abundance.

**Figure 2.**
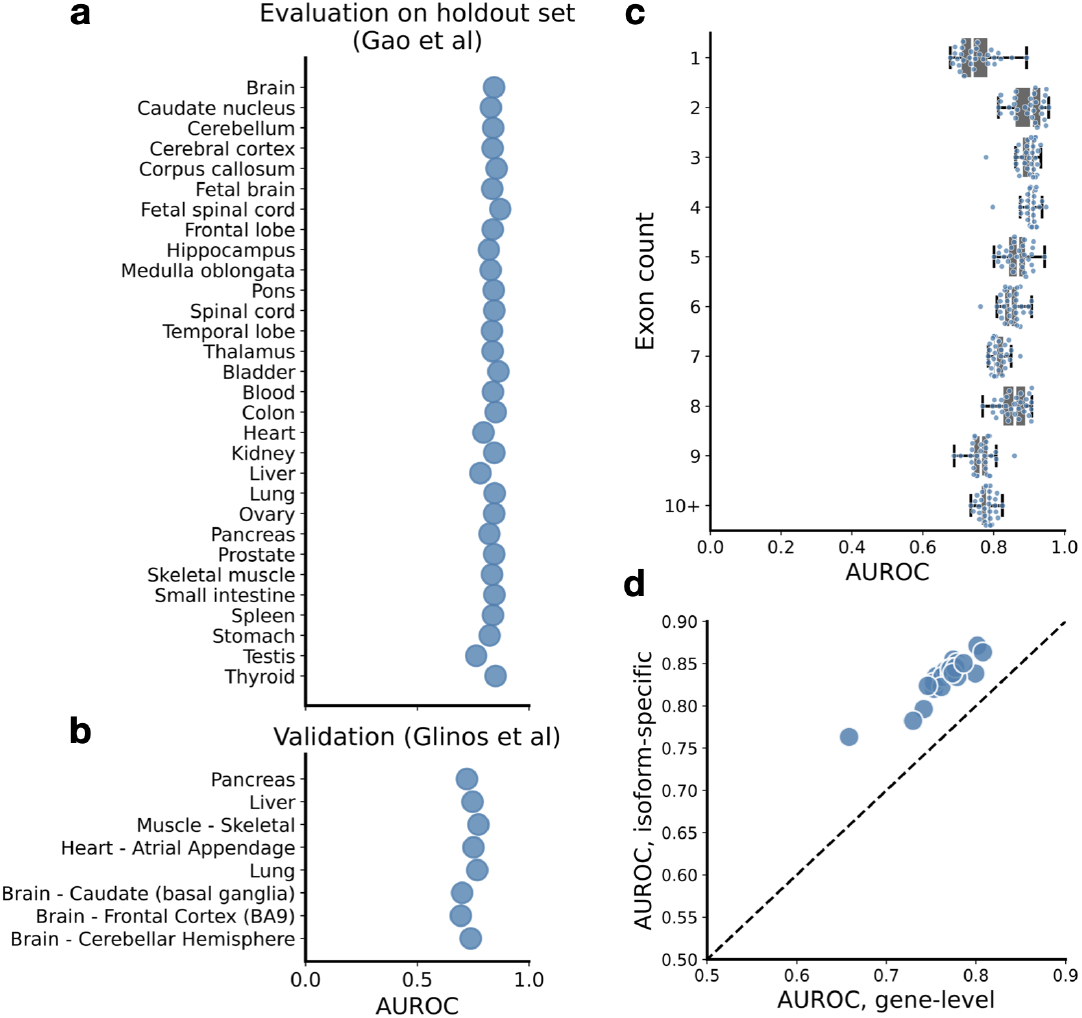
Validation of Otari on held-out and independent datasets. **a**, Performance of Otari on the Gao et al. chromosome 8 holdout test set (*n* = 5,448 canonical transcripts). Relative isoform abundances were binarized to classify high- and low-expressed transcripts. Prediction performance is shown as AUROC values (*x*-axis) across 30 tissues and brain regions (*y*-axis). Each point represents a tissue-specific model. **b**, Independent validation on an external dataset. Otari was evaluated on canonical transcripts from the Glinos et al. (*n* = 5,711) and Leung et al. (*n* = 787) validation test datasets (chromosome 8 holdout transcripts only). Glinos et al. tissues matched to Otari models are shown. **c**, Otari performance stratified by transcript complexity. Chromosome 8 transcripts from the Gato et al. holdout test set were grouped by exon count into 10 bins: nine bins for transcripts with 1–9 exons, and one bin for transcripts with ≥10 exons. AUROC distributions (*y*-axis) across tissues are shown for each bin. In all boxplots, center lines indicate the median, box limits denote the 25th and 75th percentiles, whiskers extend to 1.5× the interquartile range, and individual points represent tissue-specific values. **d**, Isoform-specific predictions outperform global gene-level comparisons. AUROC values for Otari’s isoform-level predictions (*y*-axis) were compared to gene-level predictions computed using the predicted most abundant transcript per gene (*x*-axis) in the Gao et al. holdout test set.

The generalizability of our model was supported by rigorous evaluations on independent datasets^46,47^, including 6,498 canonical transcripts from the held-out chromosome. Across matched tissues in the validation test Glinos et al. dataset^46^, Otari achieved a strong average AUROC of 0.737 (mean AUPRC = 0.730; Fig. 2b), and performed similarly well in another validation test evaluation on the Leung et al.^47^ dataset (mean AUROC = 0.657 and mean AUPRC = 0.787; Supp. Table 1) despite differences in donors, sequencing platforms, and experimental conditions from the training data. We also evaluated Otari’s ability to predict expression of novel transcripts, which are often difficult to quantify due to low abundance or limited annotation. Despite these challenges, Otari achieved a mean Pearson’s correlation of 0.368 across 196,283 novel isoforms (Supp. Fig. 4), matching its performance on lower-abundance canonical transcripts (mean Pearson’s *r* = 0.346; *p* = 0.11; two-sided independent *t*-test).

Long non-coding RNAs (lncRNAs) undergo splicing and can generate multiple isoforms, contributing to transcriptomic complexity and functional regulation^48,49^. Despite their lower conservation compared to protein-coding genes, Otari achieved comparable performance on lncRNA and protein-coding isoforms (lncRNA mean AUROC = 0.823; protein-coding mean AUROC = 0.808; Supp. Fig. 5), supporting its strong generalizability across transcript types. Notably, Otari’s performance was also sustained when aggregating predictions per gene (gene-specific Pearson’s *r* mean = 0.493 and std = 0.029; Supp. Fig. 6) and was robust to transcript complexity, as demonstrated by accurate predictions even for transcripts with 10+ exons (Fig. 2c; mean AUROC for 1-10 exons = 0.844; mean AUROC for 10+ exons = 0.781; transcript counts shown in Supp. Fig. 7).

Importantly, Otari captures transcriptomic variation that isoform-agnostic models often miss. When benchmarked on the held-out set, AUROC values based on Otari’s isoform-specific predictions consistently outperformed gene-level baselines derived from the predicted most abundant transcript per gene (Fig. 2d; mean AUROC: 0.835 vs. 0.766). These results underscore Otari’s ability to accurately model differential isoform abundance beyond gene-level expression, supporting its application in downstream isoform analyses.

### Modeling isoform-level variant effects

Genetic variants reshape transcriptomic architecture by altering splicing patterns and isoform abundance, thereby contributing to diverse disease outcomes^9,12,14,17,18,50–53^. To systematically quantify these effects, we extended Otari to predict isoform-specific consequences of single nucleotide variants (Fig. 1b). For each variant, Otari mutates the transcript graph node sequences, recomputes node features, and evaluates the reference and alternative graphs to estimate log fold changes in tissue-specific isoform abundance. Furthermore, by analyzing the graph structure, Otari identifies affected nodes, which often correspond to alternatively-spliced regions (see below). These predictions, coupled with interpretability of the underlying disrupted regulatory features, provide mechanistic insight into how variants perturb isoform profiles.

To validate Otari’s variant effect predictions, we leveraged fine-mapped expression and splicing quantitative trait loci (eQTLs and sQTLs) from GTEx v10^42^. Notably, Otari correctly inferred the direction of expression change for thousands of eQTLs across 10 tissues (Fig. 3a). Although eQTLs are defined at the gene level, aggregated isoform-level predictions achieved 87.5% accuracy for variants with strong predicted effects (normalized summed score > 64). Otari also successfully captured transcript-specific expression changes associated with sQTLs: transcripts overlapping or flanking variant-associated spliced regions showed significantly greater predicted impact than non-overlapping transcripts (Fig. 3b; max *q* = 9.49 × 10^−3^).

**Figure 3.**
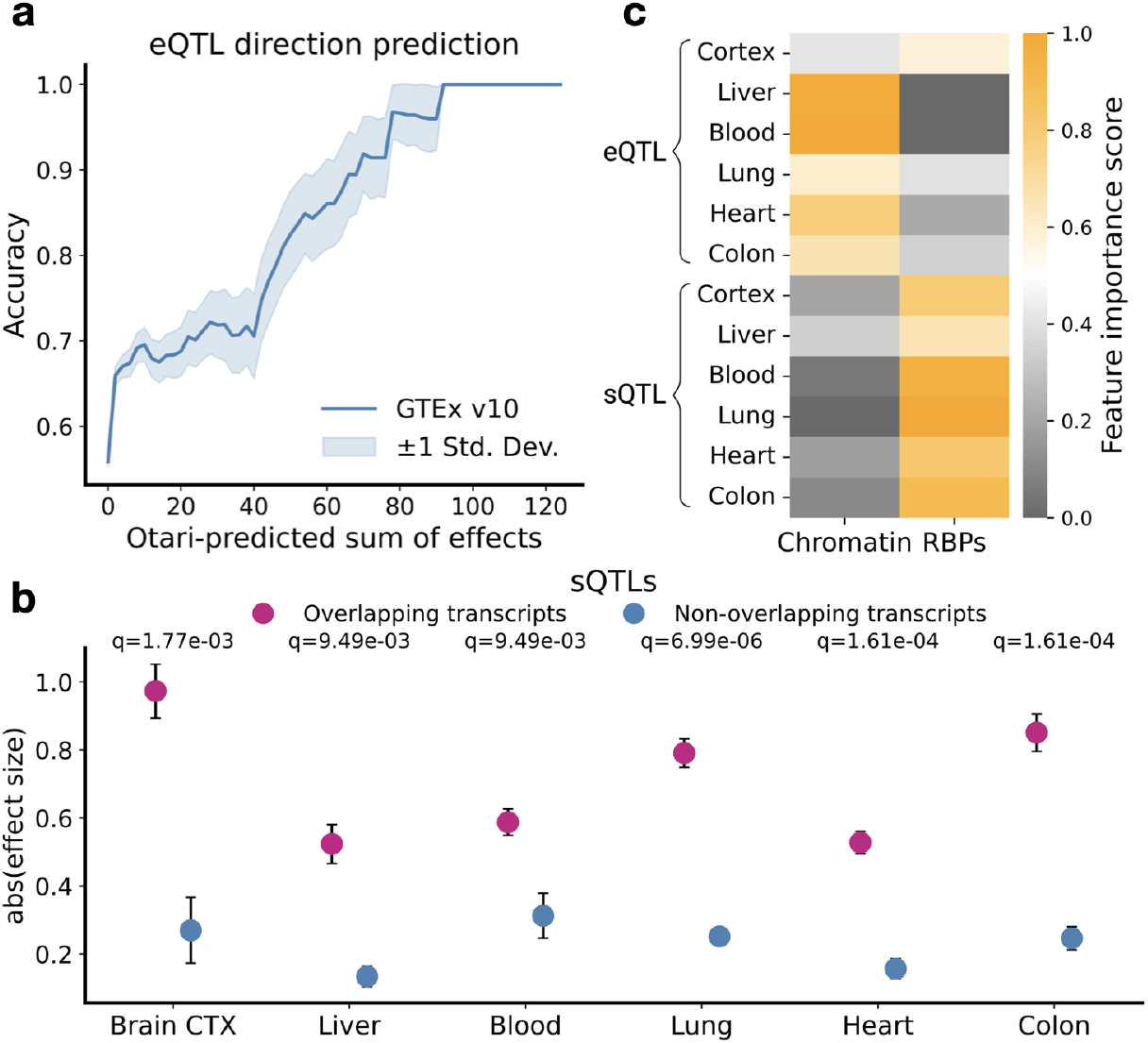
Otari captures functional effects of GTEx QTLs. **a**, Directionality prediction of fine-mapped expression quantitative trait loci (eQTLs). Variant sets from GTEx v10 across ten distinct tissues (*n* = 11,146) were analyzed. For each variant, Otari-predicted isoform-level effects were aggregated per gene and compared against the direction of the eQTL allelic fold change (aFC). Prediction accuracy (*y*-axis) was assessed across a range of summed-effect thresholds (*x*-axis). Shaded regions represent ±1 standard deviation from 1,000 bootstrap samples. **b**, Otari predicts differential abundance effects associated with fine-mapped splicing QTLs (sQTLs). sQTLs from six GTEx v10 tissues were analyzed by grouping transcripts into overlapping or non-overlapping categories based on their exon overlap with the sQTL-associated spliced region. Absolute effect sizes (*y*-axis) were compared between groups for each tissue. All statistical testing used one-sided independent *t*-tests with Benjamini-Hochberg correction. Scores were normalized relative to a background distribution (Methods). Circles denote means; error bars represent the standard error of the mean (SEM). Sample sizes (overlapping/non-overlapping) for representative tissues are: brain cortex (n = 2,947/353), liver (n = 1,795/214), whole blood (n = 4,848/580), lung (n = 6,696/909), heart (n = 4,382/512), and colon (n = 4,868/599). **c**, Functional feature burden analysis of fine-mapped QTLs. Otari-estimated variant effect scores were used to select the top impacted regulatory features for each eQTL and sQTL dataset (six tissues each, *y*-axis). Features were annotated to either chromatin-associated (Sei) or RNA-binding protein (RBP)-associated (Seqweaver) regulatory categories. Category counts were min–max normalized within each category. Color scale indicates normalized burden per category.

Otari additionally offers interpretable insights into the regulatory mechanisms driving variant effects. For each QTL, we identified the most disrupted node features, weighted by predicted effect sizes, and annotated the top features in each variant set to transcriptional (Sei) or post-transcriptional (Seqweaver) regulatory categories. Otari accurately captured the distinct regulatory signatures of QTLs: sQTL effects were primarily driven by RBP-associated features (Fig. 3c; mean Seqweaver score = 0.85; Sei = 0.15), whereas eQTL effects were mostly attributed to chromatin-based signals (Fig. 3c; Seqweaver = 0.26; Sei = 0.74). Together, these findings demonstrate that Otari enables accurate and interpretable predictions of variant-driven isoform changes, establishing a framework for isoform-level interpretation of regulatory variants.

### Disease mutations differentially impact isoforms

We next applied Otari to disease-associated functional polymorphisms (DFPs) curated in the Human Genome Mutation Database (HGMD)^40^ to assess how known pathogenic variants affect isoform profiles. Compared to control variants without disease association, DFPs showed significantly greater predicted effects on isoform abundance, highlighting strong variant-level specificity (Fig. 4a; *q* = 1.13 × 10^−95^). Otari also uncovered isoform-specific differences in variant impact within genes. For each disease variant, we identified the isoform with the strongest predicted effect. We found that many of these isoforms were annotated as non-principal based on structural, functional, and evolutionary metrics in APPRIS^54^. When we compared the predicted effects of DFPs on these highly affected isoforms versus principal transcripts, we observed significantly lower impact on the principal set (Fig. 4a; *q* = 5.34 × 10^−4^; mean effect size = 0.95 for principal, 1.41 for top-ranked), despite similar baseline expression levels (Supp. Fig. 8; mean TPM: 5.27 and 3.98). These analyses demonstrate the specificity of Otari’s predictions at both the variant and isoform levels, and suggest important functional roles for individual transcripts in disease.

**Figure 4.**
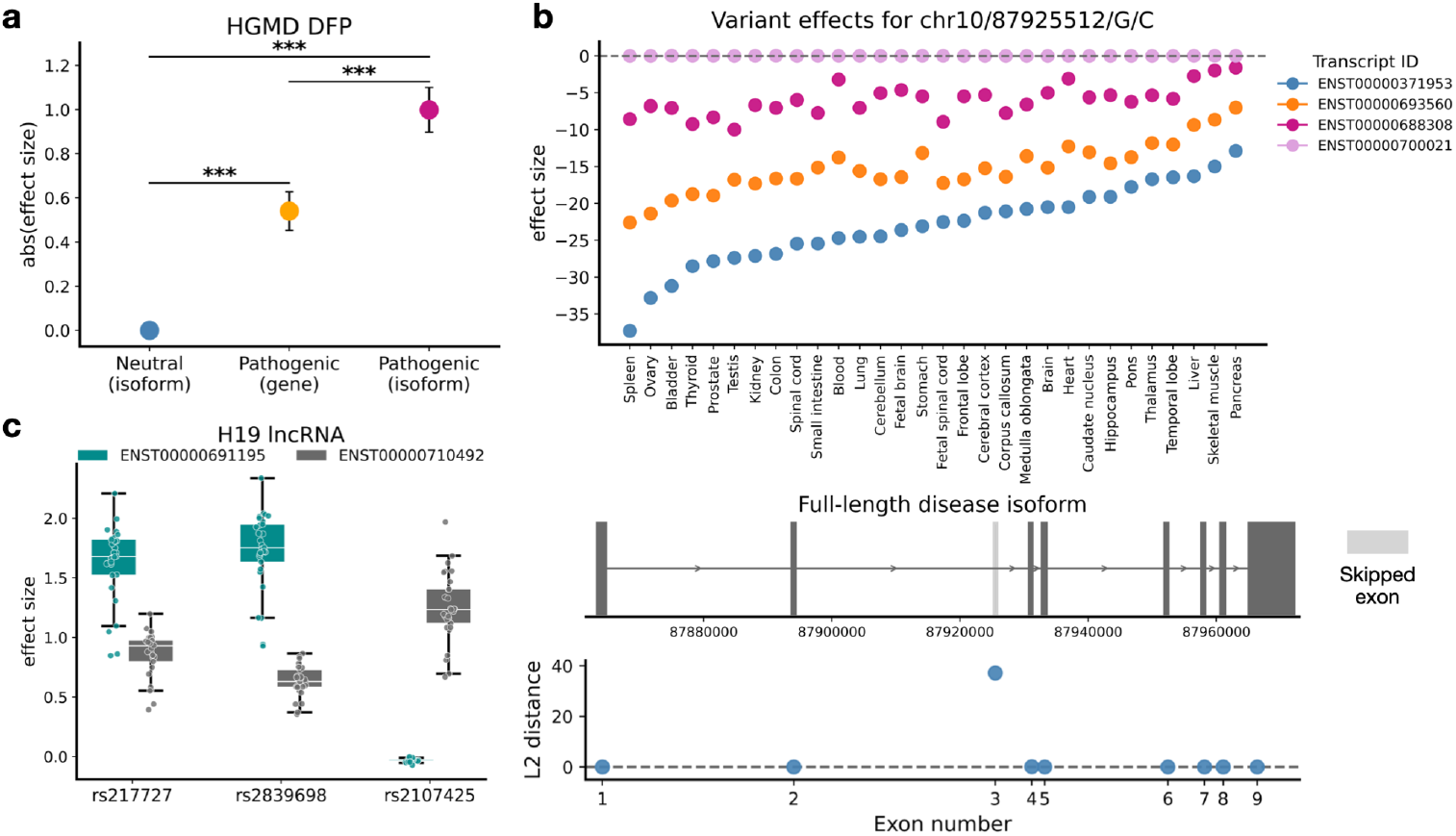
Otari predicts isoform-resolved impact of regulatory disease variants. **a**, Predicted variant effects from Otari for regulatory disease-associated functional polymorphisms (DFPs) from the Human Gene Mutation Database (HGMD) (*n* = 2,174). From right to left: distribution of DFP effects on the most impacted isoform per gene; DFP effects on the principal transcript of each gene; and a control set of neutral variants (*n* = 78,369), showing absolute effects on the most impacted isoform per gene. Effects were retained for the top impacted tissue per variant. Circles represent group means; error bars indicate the standard error of the mean (SEM). One-sided independent *t*-tests were used for hypothesis testing with Benjamini-Hochberg correction for multiple comparisons. Asterisks (***) indicate false discovery rate (FDR) < 0.01. Normalized to neutral isoform effects. **b**, Isoform-level predictions for clinical PTEN variant rs786203847. Top: effect size fold changes (*y*-axis) across 30 tissue types (*x*-axis) for four PTEN isoforms; the dashed line indicates no effect. Middle: splice structure of the affected isoform showing exon 3 skipping (lighter shade); the *x*-axis corresponds to the genomic coordinates on chromosome 10. Bottom: L2-norm distances between graph node attributes in reference and alternative sequences across the nine exons; the largest impact was observed in exon 3. **c**, Application of Otari to cancer-associated variants (rs217727, rs2839698) and a cancer-protective variant (rs2107425) in the H19 long non-coding RNA. Log fold change distributions across tissues (*y*-axis) are shown separately for each variant and two annotated isoforms. Center lines denote medians, box limits indicate 25th and 75th percentiles, whiskers span 1.5× the interquartile range, and individual tissue-specific values are plotted as points. All effect sizes were z-score normalized relative to a background distribution (Methods).

To further illustrate this, we examined a pathogenic clinical^41^ variant in PTEN, a gene that has been implicated in various neurodevelopmental disorders and cancer^55–57^. Otari predicted that variant rs786203847 selectively downregulates several annotated PTEN isoforms (Fig. 4b, top). For example, two transcripts, ENST00000371953 and ENST00000693560, showed sharp, tissue-specific predicted reduction in expression (mean effects = –23.1 and −15.2). However, Otari predicted that transcript ENST00000688308 is only moderately downregulated (mean effect = −5.99), while isoform ENST00000700021 remained unaffected (mean = 0.01; all transcripts shown in Supp. Fig. 9). Otari further identified mechanisms for these changes. Analysis of the graph nodes before and after introduction of the variant revealed that the most disrupted node corresponds to exon 3 of PTEN, consistent with previously reported exon 3 skipping for this variant^58^ (Fig. 4b, bottom). Otari also profiled the underlying disrupted regulatory signals, including loss of the donor splice site signal, increased signal of the repressive histone mark H3K27me3, and altered RNA-binding protein activity involved in splicing and transcript stability (Supp. Fig. 10).

We also demonstrate that Otari captures isoform-specific dysregulation in lncRNAs. For example, previous studies have linked three H19 lncRNA polymorphisms to cancer susceptibility, as reported in a meta-analysis of over 48,000 individuals^59^. We applied Otari to these H19-associated variants: rs217727 and rs2839698 (risk alleles) and rs2107425 (protective allele) (Fig. 4c). Among the two annotated H19 isoforms, Otari predicted that the two risk variants selectively increase expression of isoform ENST00000691195 (H19-213), consistent with reported overexpression of H19 in human cancers^59,60^. In contrast, the protective variant increased expression of a distinct isoform, ENST00000710492 (H19-227), without affecting H19-213. These findings support the hypothesis that cancer risk and susceptibility may be mediated in part by differential regulation of H19 isoforms, and propose a specific underlying mechanism for this association.

Together, these case studies highlight Otari’s ability to resolve variant-driven isoform dysregulation, including transcript-specific expression changes, splicing disruptions, and associated regulatory mechanisms that are largely undetectable in isoform-agnostic analyses.

### Isoform misregulation in autism

Transcript isoform diversity and regulation are particularly relevant to neurodevelopmental disorders, as the brain exhibits the highest levels of alternative splicing among all human tissues^8,13,61^. Otari enables direct mapping of genetic variants to isoform-specific consequences, allowing mechanistic interrogation of *cis*-regulatory variation implicated in autism and related conditions^27,62^.

We applied Otari to a large autism cohort, SPARK^43^. Whole-genome *de novo* variant (DNV) calling was performed for 3,507 autistic probands and 2,206 unaffected siblings (Methods). While overall DNV burdens were comparable between groups (Fig. 5a; mean counts: 70.92 vs. 70.44), Otari predicted significantly greater isoform dysregulation associated with proband DNVs compared to unaffected siblings in Satterstrom genes^63^ (Fig. 5b; *p* = 2.32 × 10^−7^). These effects were strongly tissue-specific, with brain regions showing significantly higher predicted disruption in isoform abundance compared to non-brain tissues (Fig. 5c; *p* = 3.97 × 10^−6^), particularly in the fetal brain, corpus callosum, and cerebellum, all of which have been associated with autism^56,62^.

**Figure 5.**
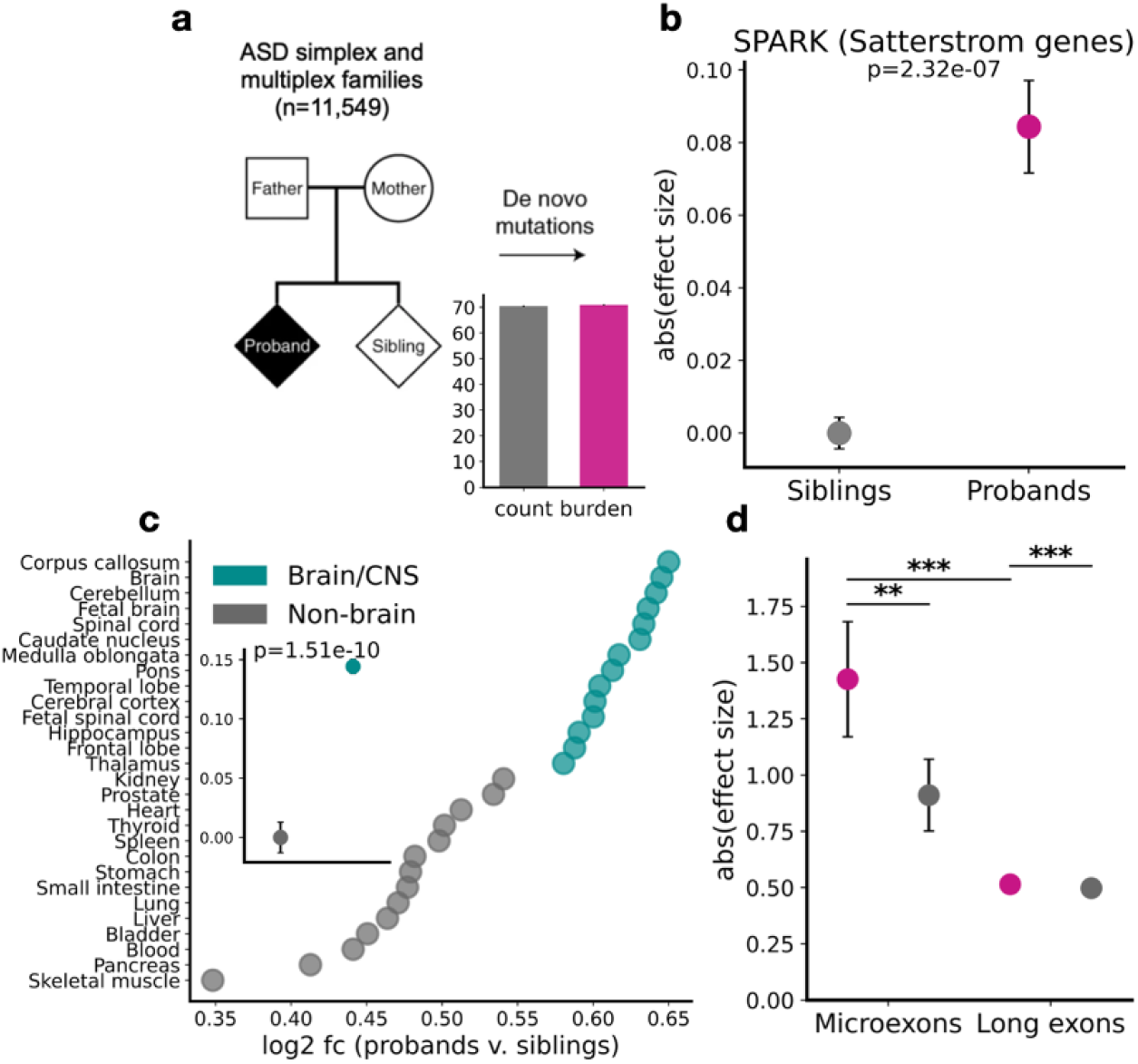
Analysis of isoform dysregulation in autism. **a**, *De novo* variants were called from whole-genome sequencing of complete trios in the SPARK cohort. Families are either simplex (one affected child) or multiplex (multiple affected children). Genome-wide *de novo* variant burden (*y*-axis) was computed for each proband and sibling. Bar heights indicate group means, with error bars representing the standard error of the mean (SEM). **b**, Otari-predicted effect size fold changes (*y*-axis) for *de novo* variants in SPARK probands (right) and siblings (left), limited to genes in the Satterstrom ASD-associated gene set. Effects were retained for the top impacted brain tissue per variant. In all analyses, circles represent means; error bars show SEM. One-sided independent *t*-tests were used for hypothesis testing, with Benjamini-Hochberg correction applied for multiple comparisons. All effect sizes were z-score normalized relative to a background distribution (Methods). **c**, For each tissue, the log2 fold change (fc) of the mean absolute Otari score for probands compared to siblings is shown (*x*-axis). The inset compares the distribution of log2 fc between brain/CNS tissues and non-brain tissues. **d**, Microexon misregulation is associated with *de novo* variants in SPARK. Otari-predicted effects were computed for *de novo* variants mapping to brain-expressed genes (*n* = 57,929 variants in probands, *n* = 35,642 variants in siblings). Effect sizes (mean ± SEM) are shown for variants located nearby splice sites for microexons (3-27 nt; *n* = 650/355 transcripts) and long exons (>27 nt; *n* = 63,786/38,794 transcripts). Effects were retained for the top impacted brain tissue per variant. Significance stars are marked as follows: one star (*) indicates FDR < 0.1, two stars (**) FDR < 0.05, and three stars (***) FDR < 0.01.

To investigate some of the mechanisms underlying these effects, we examined microexons, which are highly-conserved, neuron-specific short exons (3–27 nt) previously implicated in autism and other neurodevelopmental disorders^64–67^. We classified all brain-expressed exons as either microexons or long exons (>27 nt) and assessed the impact of nearby *de novo* variants. Notably, transcripts containing microexons near proband variants showed significantly greater dysregulation than those with nearby long exons (Fig. 5d; *q* = 6.0 × 10^−4^). Proband variants near microexons also drove significantly stronger expression changes than sibling variants near microexons (*q* = 0.04), suggesting altered microexon inclusion in autistic individuals. Together with prior small-scale experimental studies^64–66^, our findings from analysis of a large cohort support variant-driven microexon misregulation as a prevalent feature of autism pathophysiology.

## Discussion

Otari is a sequence-based model capable of resolving isoform-level regulatory and expression changes at scale. This attention-based graph deep learning framework offers a powerful approach for transcriptome-wide, isoform-level variant effect analysis, driving insights into the regulation of isoform usage in various biological contexts. Unlike prior models that focus primarily on predicting individual splice sites or RBP binding, Otari learns rich, structure-aware representations of full transcripts, capturing the complex regulatory code underlying differential isoform abundance. Trained on long-read transcriptomic data spanning diverse tissue types, Otari provides high-resolution predictions across canonical and novel isoforms, generalizes to complex multi-exonic structures, and drives isoform-resolved variant interpretation.

Otari also provides interpretable insights into how genetic variation alters isoform usage. Across large-scale mutation datasets, Otari enabled high-specificity isoform-level variant effect predictions. For example, Otari predicted that a regulatory clinical variant in PTEN led to minimal to modest downregulation of some transcripts, but stronger, more tissue-specific effects on alternative transcripts, along with exon skipping and regulatory disruptions involving H3K27me3 and RNA-binding proteins. In H19, Otari revealed that cancer risk and protective variants modulate the expression of distinct lncRNA isoforms, demonstrating its capacity to resolve functional mechanisms even in low-conservation regions. These findings open a new avenue for systematic studies of lncRNA isoforms in future research.

Applying Otari to a large autism cohort further demonstrated its potential for mechanistic discovery. Despite similar genome-wide variant burdens, we observed significant brain-specific isoform dysregulation in autistic probands compared to unaffected siblings, particularly in brain regions previously implicated in autism. Otari also captured altered microexon usage in probands compared to unaffected siblings based on analysis of a large cohort, further supporting the role of this mechanism in autism pathophysiology previously reported in small-scale experimental studies. These findings point to a broader role for isoform misregulation in neurodevelopmental disorders. Otari could be applied to study other conditions characterized by splicing abnormalities, such as schizophrenia, ALS, and Alzheimer’s disease. Notably, Otari’s demonstrated ability to interpret regulatory variants at the isoform level, which are challenging to study experimentally and at scale, offers a promising path for isoform-resolved variant interpretation in a wide variety of context-specific studies.

In future work, integrating single-cell long-read sequencing will be critical for capturing isoform regulation at cell-type resolution. While Otari models tissue-level regulation, resolving cell-type-specific regulation is essential in heterogeneous tissues such as the brain. Incorporating these data could also improve detection and analysis of rare, novel, or cell-type-specific isoforms that may play important functional roles. Isoform regulation also varies across developmental stages and conditions. As more data become available, modeling these context-dependent regulatory dynamics will help reveal how isoform usage is altered across diverse biological and disease contexts. Another challenge lies in translating isoform-level effects to functional outcomes. While Otari provides high-specificity predictions of changes in transcript isoform abundance, linking these effects to downstream consequences, such as protein abundance, function, localization, or stability, remains non-trivial, especially for novel isoforms. This underscores the need for integration of multi-modal data types to drive prediction across multiple layers of regulation.

Despite these challenges, Otari establishes a foundation for transcriptome-wide, isoform-level studies, enabling a new level of interpretability and precision in large-scale analyses of isoform regulation, diversity, and variant impact. Future work with focus on experimental validation will be critical for translating these insights into isoform-targeted therapeutic strategies.

## Online Methods

### Ethics

We received approval to access and analyze de-identified genetic and phenotypic data from the SPARK autism cohort from SFARI Base and the Princeton University IRB Committee in the Office of Research Integrity.

### Data

#### Training and testing

A total of 135,638 unique canonical transcripts meeting filtering criteria were retrieved from the Gao et al.^45^ dataset. Gao et al. provide robust quantification of transcript isoforms using full-length long-read Nanopore sequencing across 30 distinct human tissues and brain regions. They report a total of 623 million ONT 1D cDNA reads (reads per tissue range from 13.2 to 30.6 million). Each sample included tissues pooled from between 2 to 64 individuals (see Table S5 in Gao et al.). Only transcripts annotated as full splice matches in the GENCODE v47 human reference catalog (GRCh38.p14, basic annotation)^68,69^ were used for training, validation, and evaluation. Transcripts located on the X, Y, and mitochondrial (M) chromosomes were excluded from all training, validation, testing, and downstream analyses. The test set consisted of all transcripts from chromosome 8 annotated as protein-coding or lncRNAs (*n* = 5,448), and all other transcripts from other autosomes were used for training and validation. Isoform abundance values (in TPM) were log_2_-transformed with a pseudocount of 0.01 across all datasets.

#### Validation

Model validation was conducted using three independent transcript sets: novel isoforms from Gao et al., and canonical isoforms from the Glinos et al.^46^ and Leung et al.^47^ datasets. The Gao et al. set comprised 196,283 unannotated novel transcripts with annotated splice sites, profiled across 30 tissues. The Glinos et al. validation test set included 5,711 transcripts (protein-coding and lncRNA only) from chromosome 8, profiled via Nanopore sequencing across multiple tissues. The Leung et al. validation test set consisted of 787 chromosome 8 transcripts (protein-coding and lncRNA only) profiled in fetal and adult cortex using both Nanopore and PacBio platforms. For each dataset, transcript expression was reported in TPM (transcripts per million), and values were averaged across samples and/or replicates within tissues. Only transcripts with full splice match annotations were included. Transcripts were annotated as protein-coding or lncRNA based on the GENCODE v47 reference to assess gene biotype distribution.

### Graph construction

#### Graph structure

We constructed a directed graph for each transcript across all datasets, where nodes represent the sequence context surrounding the 3′ and 5′ splice sites of each exon, and edges represent the sequential ordering of exons based on the GENCODE reference annotation. In the absence of node features, the resulting gene structure resembles a standard splice graph. Node sequences were generated in 5′ to 3′ order. For transcripts on the positive strand, nodes were ordered by increasing genomic position; for those on the negative strand, nodes were ordered by decreasing position.

#### Node attributes

To create a rich feature representation for each node, we constructed a concatenated vector of biological signals using three deep learning models applied to the DNA sequence context around each splice site. For splicing signals, we used a custom model named ConvSplice (described below) to score the likelihood that each site represents a splice donor or acceptor. For post-transcriptional regulation, we generated Seqweaver^36^ predictions as part of the node embedding vector. Seqweaver provides predicted binding affinity scores for >100 RNA-binding proteins (RBPs), including regulators of splicing, stability, localization, and translation, based on 217 Cross-Linking and Immunoprecipitation sequencing (CLIP-seq) datasets. For each splice site, we generated predictions across an extended 1 kb window using eight 50-nt bins spanning from 200 bp upstream to 200 bp downstream. Finally, we used the Sei model^26^ to predict chromatin-associated regulatory features as part of the node embeddings, including histone marks, transcription factors, and DNAse sensitivity. We retained the 10,062 *cis*-regulatory peak scores for histone marks (e.g., H3K27ac, H3K4me3), which have been previously linked to splicing and isoform control^23^. Predictions were obtained using a 4 kb sequence window centered on each splice site. These outputs were concatenated to form a final node feature vector of dimension 23,600, structured as follows: 3’ SpliceConv(2), 3’ Seqweaver(1,736), 3’ Sei(10,062), 5’ SpliceConv(2), 5’ Seqweaver(1,736), and 5’ Sei(10,062). The numbers in the parentheses indicate the feature dimension for each component. All genomic sequences were extracted from the GRCh38 human reference genome.

#### ConvSplice model

ConvSplice is a deep learning model developed to predict splice donor and acceptor sites directly from DNA sequence. It was trained using principal transcripts from GENCODE v40^69^ annotated by APPRIS^54^. Positive examples included known donor (5′) and acceptor (3′) splice sites; all other positions in the transcripts were treated as negatives. To evaluate the model’s performance, we used chromosomes 1, 3, 5, 7, and 9 as the test set, with the remaining chromosomes used for training and validation.

The ConvSplice architecture builds upon the framework introduced by SpliceAI^30^ but incorporates several key improvements. The model takes as input a 20kb sequence window centered on the position of interest (10kb on each side). Each nucleotide position in the input sequence is one-hot encoded as a 4-dimensional vector representing A, C, G, and T. The model outputs three scores for each position, representing the probability of that position being a splice acceptor, splice donor, or neither. The architecture consists of 20 dilated convolution layers with residual connections applied every four layers. Each layer includes batch normalization and ReLU activation functions followed by convolution operations. This deeper architecture with a larger sequence context enables the model to capture more complex splicing patterns compared to previous approaches.

For model training, we randomly split the training data into 80% training and 20% validation sets. The model was trained using the Adam optimizer with an initial learning rate of 0.0001. We utilized the ReduceLROnPlateau (PyTorch v2.2.2)^70^ learning rate scheduler that monitored validation loss to adaptively adjust the learning rate during training. To ensure robust predictions, we trained 5 independent models using different random seeds and used the average of their predictions as the final output. Performance compared to SpliceAI can be found in Supp. Fig. 1b.

### MPNN training

#### Overview

The model was trained to predict relative isoform abundances across 30 distinct tissue types and brain regions by mapping a learned graph embedding to the target abundance values. Training was performed on mini-batches of transcripts, each encoded as a directed graph where nodes represent the sequence context surrounding the 3’ and 5’ splice sites, and edges represent exon connectivity as annotated in GENCODE v47. The model output is continuous, predicting log2-transformed relative isoform abundances. Chromosome 8 was held out as a test set, while all other autosomes were used for training and validation (X, Y, and M were excluded). The implementation was built with PyTorch Geometric v2.5.2^71^ and PyTorch Lightning v2.3.0, with training tracked using Weights and Biases v0.17.0.

#### Architecture search

To optimize model performance, an architecture search was conducted to evaluate different configurations. Candidate models were trained with either 2, 3, or 4 stacked attention-based message-passing blocks, and comparisons were made between models using graph attention networks (GAT) and graph convolutional networks (GCN) to assess the added benefit of attention mechanisms. Global pooling strategies, including add, mean, and max, were benchmarked, along with the effects of concatenating versus averaging attention head outputs. Residual connections were also tested to evaluate whether propagating information from previous layers improved performance. All models were trained for 50 epochs during architecture benchmarking.

The final selected architecture consisted of three sequential components: (1) three blocks of graph attention layers with residual connections; (2) a global pooling operation using maximum aggregation; and (3) a two-layer fully connected output head. Each attention block included two GAT layers followed by batch normalization, ReLU activation, and dropout. Residual connections were applied in the second and third attention blocks to enhance information flow across layers. After attention-based message passing, node embeddings were pooled to produce a graph-level embedding, which was then processed by a feed-forward head consisting of linear layers, dropout, and non-linear activations to generate a prediction for each transcript across the 30 tissues.

#### Hyperparameter sweeps

Hyperparameter sweeps were conducted using Bayesian optimization via the Weights and Biases sweep utility. The search explored a range of learning rates (from 0.001 to 0.000001), hidden channel dimensions (180, 220, 320, 440, 512, 840, 932), dropout rates (0.1 to 0.5), batch sizes (32 and 64), and attention head counts (2, 4, and 8). For each sweep configuration, 80% of transcripts (excluding those on chromosome 8) were used for training and 20% for validation, with a maximum of 15 training epochs.

The final model was trained using the selected hyperparameters: learning rate of 0.0002646, hidden channel size of 512, dropout rate of 0.5, batch size of 64, and 2 attention heads. Training was performed on a NVIDIA V100 GPU and took approximately 8 hours. To enhance the model’s learning on challenging examples, hard example mining was incorporated during training. Specifically, a weighted hard loss was computed using the TripletMarginLoss function in PyTorch v2.2.2 for each batch, and this was combined with the mean squared error loss to form the total training objective:

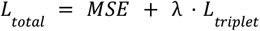

Where:

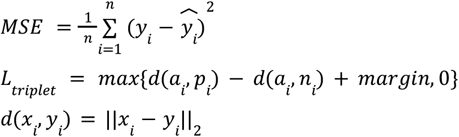

In this equation, MSE represents the mean squared error between the predicted value 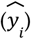 and the experimental measured value (*y*_*i*_). The triplet loss term *L_triplet* forces the model to learn the patterns where similar transcripts are closer together and dissimilar ones are further apart. We set *λ* = 0.25 and *margin* = 1.0. For each training batch, anchor (*a*_*i*_), positive (*p*_*i*_), and negative (*n*_*i*_) examples were selected based on transcript expression patterns, with *L*2 norm as the distance metric *d*.

### Model performance and evaluation

#### Abundance binarization

To evaluate the model’s ability to distinguish between low and high expressed transcripts in each tissue, we binarized transcript abundances based on percentile thresholds. Specifically, transcripts with expression values above the 70th percentile within a tissue were labeled as high abundance (positive examples), while those below the 30th percentile were labeled as low abundance (negative examples) using ground truth data. These percentile thresholds were tissue-specific and changed according to the expression distribution within each tissue. Predicted transcript abundances were ranked and scaled to a (0,1] interval by normalizing against the total number of transcripts. Model performance on this binarization task was assessed using the chromosome 8 holdout set from the Gao et al. dataset, along with validation on chromosome 8 transcripts from the Glinos et al. and Leung et al. validation datasets. Tissue-specific thresholds were computed separately for each dataset, and tissues with available Otari models were kept for evaluation. This included eight tissues in Glinos et al. and two tissues in Leung et al.

#### Evaluation on novel isoforms

The model was also evaluated on its ability to predict abundance levels of novel unannotated isoforms. A total of 196,283 novel isoforms from the Gao et al. dataset were included in this analysis, with transcript graphs generated using available exon annotations from Gao et al.. Predictions for novel isoforms were compared to measured abundances from the study, and Pearson correlation coefficients were computed for each tissue. In comparison, we assessed performance on lower-abundance canonical transcripts, defined as those with expression levels below the 80th percentile in each tissue.

#### Global-versus isoform-level predictions

To compare gene- and isoform-level performance, we computed AUROC scores at both levels. We compared isoform-specific performance to a baseline that assigns each gene the maximum predicted expression among its isoforms (max-predicted isoform). These gene-level maxima were evaluated against ground-truth binarized isoform expression across all transcripts. Isoform-specific AUROC scores were computed as described previously, by comparing predicted and true binarized abundances at the individual transcript level.

#### Transcript complexity

We stratified transcripts on chromosome 8 from the Gao et al. holdout test set into ten bins based on exon count, including bins for transcripts with 1 through 9 exons and a final category for those with 10 or more exons. For each category, we reported the range of tissue-specific AUROC scores to understand how prediction performance is affected by exon count.

#### Performance within genes

We computed the Pearson correlation between predicted and observed isoform abundances for all genes with at least two isoforms in the Gao et al. holdout test set. This intra-gene analysis was performed across 2,649 transcripts with available data.

### Variant effects on isoforms

#### Variant effect prediction

To quantify how variants impact differential isoform abundance, we extended the model to incorporate *cis*-regulatory variant effects. This was done by introducing sequence mutations and updating all node features using ConvSplice, Seqweaver, and Sei, allowing the model to capture functional disruptions affecting splicing and abundance regulation. Abundance predictions were made for both reference and alternative graphs, and isoform-level variant effects were defined as the log-fold change between the alternative and reference predictions per tissue: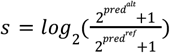.

#### Evaluation on HGMD variants

We evaluated the model on clinical variants from HGMD^40^ (release 2024.1), focusing on disease-associated functional polymorphisms (DFPs). DFPs are primarily noncoding variants found to have a statistically significant association with a clinical phenotype such as gene expression or splicing. We retained only autosomal noncoding SNPs within gene bodies or 2 kb of the TSS (*n* = 2,174). Variant effect scores were computed for each isoform by comparing predictions on reference and mutated graphs, and summarized per gene by taking the maximum absolute score across tissues and isoforms. Principal transcripts were defined using APPRIS “PRINCIPAL:1” annotations, and only the most affected principal transcript per gene was retained. For benchmarking, we compared pathogenic variant effect sizes to a control distribution, comprising the combined distribution of benign ClinVar variants and all SPARK sibling *de novo* variants passing filtering criteria (*n* = 78,369 variants, *n* = 304,851 transcript-variant combinations). All effect sizes were also z-score normalized to this background distribution. Statistical comparisons between groups were conducted using one-tailed independent *t*-tests with Benjamini-Hochberg correction.

#### Variant case studies

We also conducted case studies on select clinical variants^41,59^. Isoform variant effect scores were scaled to the background distribution described above, and L2 norm distances were calculated between reference and alternative node attribute vectors to identify the most impacted graph nodes. Features with the highest predicted change in these nodes were reported to highlight the underlying regulatory mechanisms.

### Fine-mapped QTLs

#### Datasets

We analyzed fine-mapped eQTLs and sQTLs identified using SuSiE across 49 GTEx v10 tissues^42^. Only single-nucleotide variants within genic regions or within 2 kb of the TSS were retained. Unless otherwise specified, analyses focused on six representative tissues with available models: brain cortex, liver, whole blood, lung, heart, and colon. All statistical tests were one-tailed independent *t*-tests, corrected for multiple comparisons using the Benjamini–Hochberg method.

#### eQTL directionality

To assess directionality, we compared Otari’s aggregated transcript-level predicted effects (normalized to the background distribution described above) to the known allelic fold-change (aFC) direction of *n* = 11,146 tissue-specific eQTLs. For normalized summed-score thresholds ranging from 0 to 125 (step size = 2), we summed Otari’s predicted isoform effects for each unique variant, gene, and tissue combination (for the matched tissue model scores) and filtered variants according to the threshold. The accuracy of alignment between predicted and observed aFC directions was computed at each threshold, with variability estimated via 1,000 bootstrap replicates. This analysis spanned ten tissues and brain regions, including brain cortex, hippocampus, cerebellum, liver, whole blood, pancreas, heart, lung, kidney, and colon.

#### Transcript-specific sQTL effects

For each sQTL, transcripts were split into overlapping (exons intersecting or bordering the sQTL spliced region) and non-overlapping sets. We computed the predicted effect on transcript abundance for each set within the relevant matched tissue, enabling a direct comparison of sQTL impact across isoforms.

#### Estimating tissue-specific burden

For each QTL-overlapping isoform, we identified the most perturbed graph node, defined by the largest L2 norm distance between reference and alternative node attributes, and extracted the top 10 disrupted features. Feature contributions were weighted by the sum of predicted absolute QTL effects in the matched tissue, and normalized by variant count to enable cross-tissue comparison. The top 750 features in each tissue variant set were retained and annotated to functional categories based on their model: Sei (transcriptional) or Seqweaver (post-transcriptional). ConvSplice-derived features were excluded due to the small feature space. Variants shared between the eQTL and sQTL tissue-matched sets were removed in this analysis, and min–max normalization was applied to Sei and Seqweaver feature counts separately across tissues.

### Isoform dysregulation in autism spectrum disorder

#### Datasets

The SPARK^43^ cohort provides comprehensive phenotypic, clinical, and genetic data from autistic individuals and their families. We analyzed the genetic data of 3,507 probands and 2,206 non-autistic siblings from SPARK with available trios (mother, father, child). In total, we identified 227,544 *de novo* variants (DNVs) in SPARK probands and 142,098 in siblings. Within Satterstrom genes, we compared 1,088 *de novo* variants from probands with 627 *de novo* variants from siblings. We excluded ‘testis’ and ‘ovary’ tissues from subsequent analyses due to known sex-related biases in autism^72^.

#### De novo variant calling

DNVs were identified using a pipeline from prior work^73^ and applied to whole genome sequence (WGS) data from Variant Call Format (VCF) files provided by the SPARK consortium. Briefly, the pipeline, which was optimized for WGS, uses HAT^74^ alongside variant calls from GATK HaplotypeCaller^75^ to detect variants present in the child (genotype 0/1 or 1/1) but absent in both parents (genotype 0/0). Variants were filtered based on quality metrics including read depth, genotype quality score, and genomic location. Regions with recent repeats, low complexity, or centromeric content were excluded. We used default thresholds of minimum depth = 10 and genotype quality (GQ) ≥ 20. Individuals with abnormally high DNV counts (>3 standard deviations above the mean across trios) were removed, and non-singleton DNVs (those found in more than one family) were excluded. This approach yielded an average of 70.92 DNVs per proband (s.d. = 15.07) and 70.44 per sibling (s.d. = 14.77) in SPARK.

#### Assessing average and tissue-specific impact

We applied additional filtering to exclude indels and variants on chromosomes X, Y, and M. Otari was applied to the remaining SNPs from both probands and siblings. DNVs in SPARK were restricted to Satterstrom genes (genic regions or within 2kb of TSS). For each DNV in SPARK, we computed the maximum absolute effect across brain tissues to compare effect sizes between probands and siblings. To assess tissue specificity, we computed the mean Otari-predicted effect per tissue in probands and siblings and then calculated the log fold change for each tissue. We compared the mean log fold change across brain tissues to that of non-brain tissues using a one-sided independent *t*-test. Variant effect sizes were z-score normalized to the combined distribution of benign ClinVar variants and all SPARK sibling *de novo* variants passing filtering criteria (*n* = 78,369 variants, *n* = 304,851 transcript-variant combinations).

#### Microexons analysis

We applied Otari to *de novo* variants identified in SPARK probands (*n* = 57,929 DNVs) and siblings (*n* = 35,642 DNVs), restricting the analysis to variants mapping to brain-expressed genes (*n* = 13,774 genes). Variants were located within genic regions or within 2 kb of the transcription start site. Exons were classified as either microexons (3–27 nt) or long exons (>27 nt). We flagged proband and sibling variants within 650 nt of microexon splice sites (*n* = 650 and 355 transcripts, respectively), as well as proband and sibling variants within 650 nt of long exon splice sites (*n* = 63,786 and 38,794 transcripts, respectively). Otari-predicted effects were compared between these sets; effects were retained for the top impacted brain tissue per variant. One-sided independent *t*-tests were used for hypothesis testing, with Benjamini-Hochberg correction applied for multiple comparisons.

## Acknowledgements

We are grateful to all the families in SPARK, the SPARK clinical sites and SPARK staff. We appreciate obtaining access to the SPARK phenotypic and genetic datasets on SFARI Base. Approved researchers can obtain the SPARK population data set described in this study by applying at https://base.sfari.org. This work was conducted utilizing the computing resources, supported by the Scientific Computing Core, at the Flatiron Institute. This work was supported by funding from the NIH NIGMS no. R01GM071966 (O.G.T.), Simons Foundation grant 395506 (O.G.T.), and the NIH NHGRI training grant T32HG003284 (A.L.).

## Data Availability

Training, validation, and testing data were obtained from Gao *et al*., Glinos *et al*., and Leung *et al*.^45–47^. eQTL and sQTL datasets were obtained from the GTEx v10 release^42^. Human regulatory disease mutations were obtained from HGMD (2024.1 release)^40^. Pathogenic and benign human regulatory variants were obtained from ClinVar^41^.

## Code Availability

The Otari framework code is available at https://github.com/FunctionLab/otari and the model and associated data files can be downloaded by following the instructions in the GitHub repository. The code for the manuscript results is available at https://github.com/FunctionLab/otari-manuscript.

