## Supplemental Figures for "Variant-resolved prediction of context-specific isoform variation with a graph-based attention model"

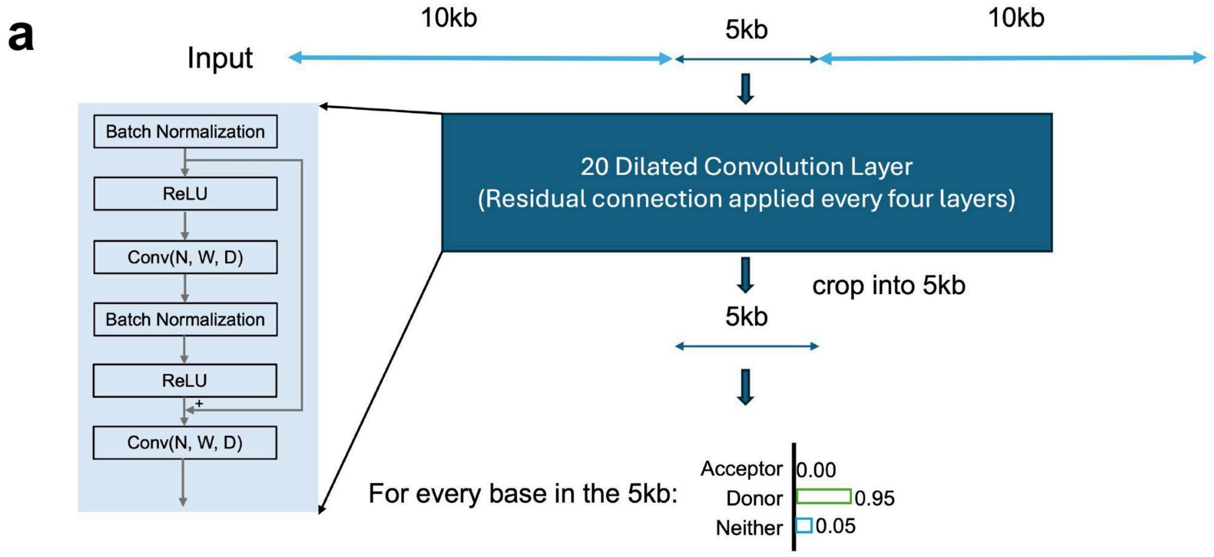


**b**

| Metric | Task | ConvSplice (20k) | ConvSplice (10k) | SpliceAI |
|---|---|---|---|---|
| **AUPRC** | Donor prediction | **0.98** | 0.97 | 0.97 |
| | Acceptor prediction | **0.98** | 0.97 | 0.97 |
| **Top-*k* Accuracy** | Donor prediction | **0.94** | 0.93 | 0.93 |
| | Acceptor prediction | **0.94** | 0.92 | 0.92 |

**Supplementary Fig. 1 | ConvSplice model. a**, ConvSplice is a deep learning convolution-based framework for splice site strength prediction. The model takes as input a 20kb sequence window centered on the position of interest (10kb on each side). Each nucleotide position in the input sequence is one-hot encoded as a 4-dimensional vector representing A, C, G, and T. The model outputs three scores for each position, representing the probability of that position being a splice acceptor, splice donor, or neither. The architecture consists of 20 dilated convolution layers with residual connections applied every four layers. Each layer includes batch normalization and ReLU activation functions followed by convolution operations. **b**, Performance comparison between ConvSplice versions and SpliceAI. ConvSplice with 20kb context outperforms both ConvSplice with 10kb context and SpliceAI across all metrics, achieving higher AUPRC (0.98 for both donor and acceptor prediction) and Top-k Accuracy (0.94 for both donor and acceptor prediction). Top-k Accuracy measures the percentage of cases where the correct splice site is among the k highest-scoring positions predicted by the model.

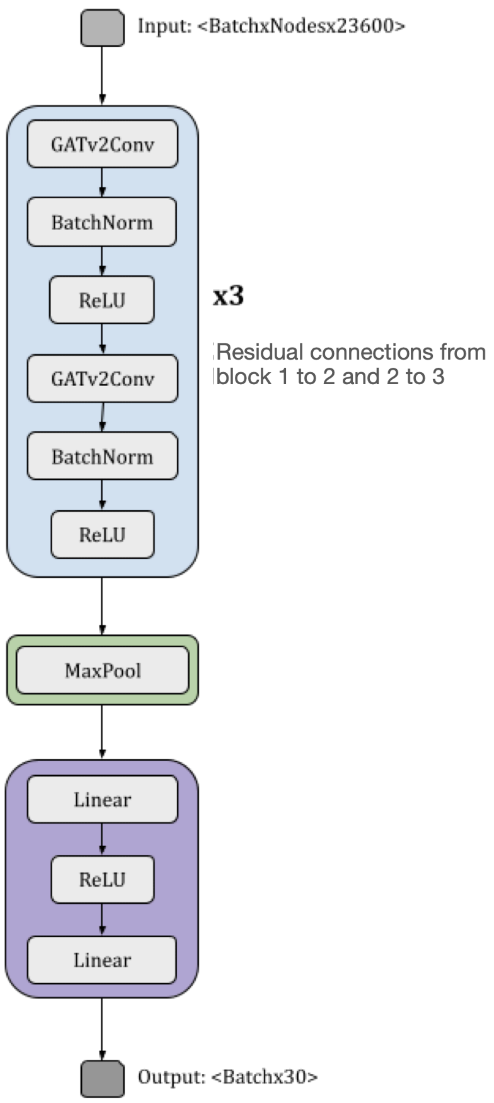


**Supplementary Fig. 2 | Otari model architecture.** Otari comprises three main modules: stacked graph attention blocks with residual connections and attention (blue), a pooling layer (green), and two fully connected output layers (purple). Each graph attention block contains two layers with batch normalization, ReLU activation, and dropout. Residual connections in the second and third blocks enhance feature propagation. Node embeddings are globally pooled using max aggregation and passed through a feed-forward head to predict isoform abundances across 30 tissues.

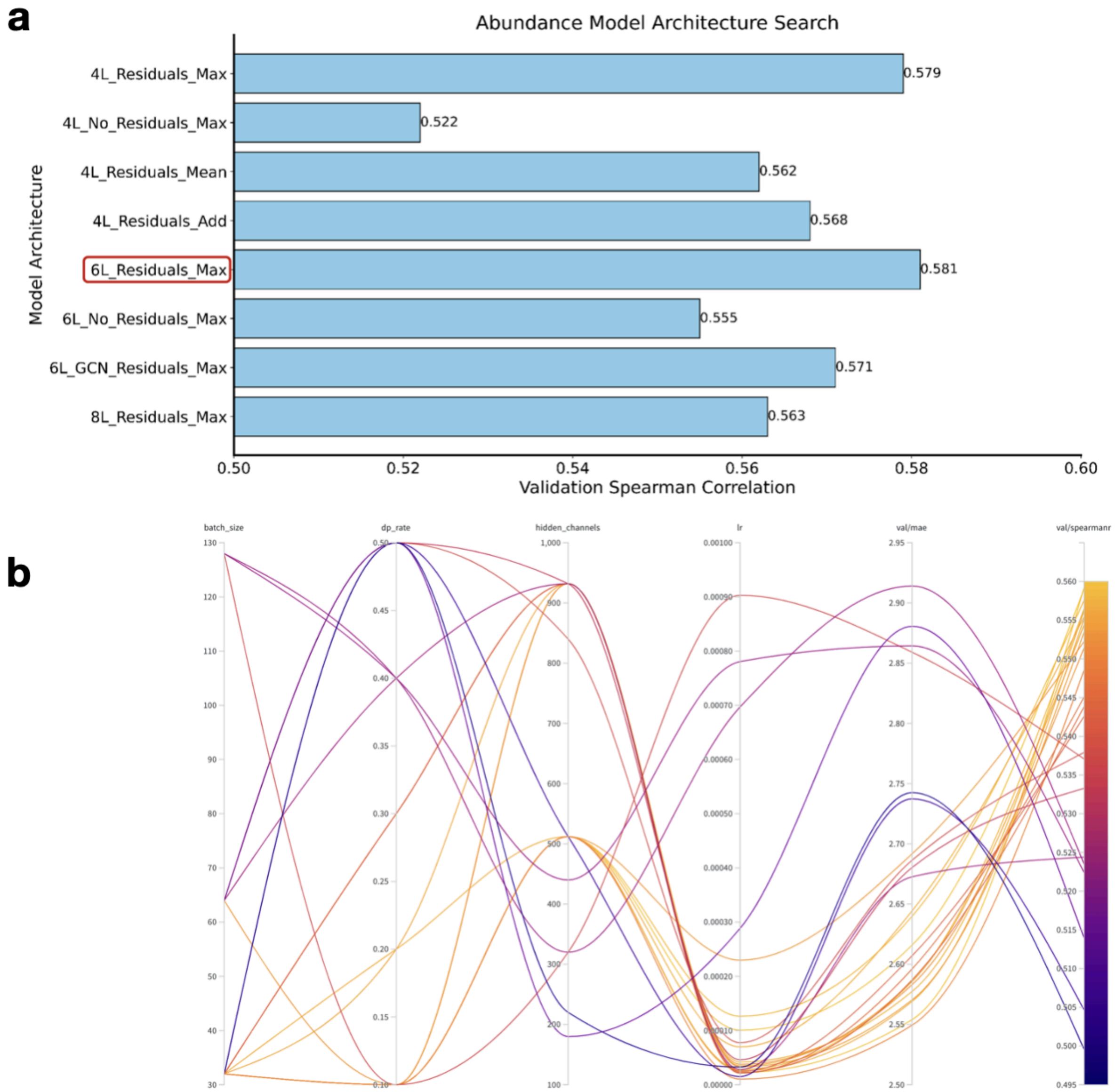


**Supplementary Fig. 3 | Model architecture and hyperparameter optimization. a**, Candidate architectures were evaluated based on validation set Spearman correlation, including models with 2–4 graph attention blocks, graph convolution (GCN), pooling strategies, attention head configurations, and residual connections. Selected model is marked in red. **b**, Hyperparameter search using Bayesian optimization across learning rate, hidden channels, dropout, batch size, and number of attention heads. Color bar indicates validation set Spearman correlation.

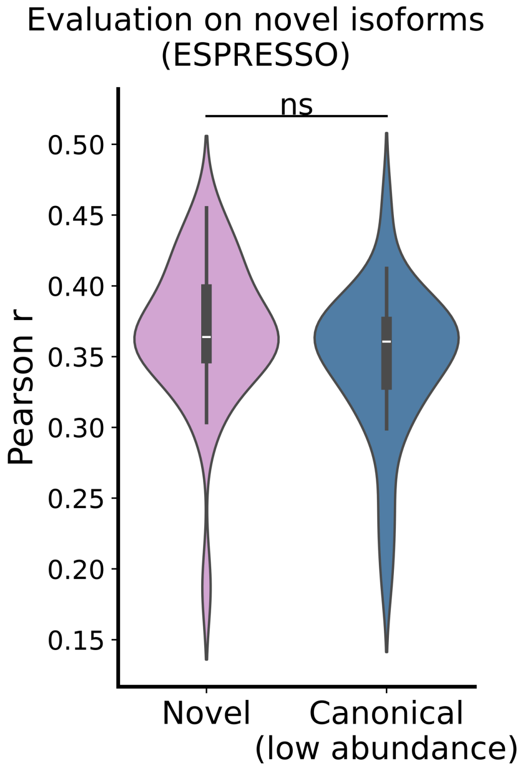


**Supplementary Fig. 4 | Evaluation of Otari on novel isoforms.** Distribution of tissue-specific Pearson correlations (*y*-axis) for novel (unannotated) isoforms from Gao et al. compared with lower-abundance canonical isoforms, defined as expression below the 80th percentile per tissue, from the Gao et al. holdout chromosome 8 test set. Boxplot center lines indicate the median, box limits denote the 25th and 75th percentiles, and whiskers extend to 1.5× the interquartile range. Violin plots additionally show the full distribution of the data. Two-sided independent *t*-test: $p$ = 0.11 (not significant). Sample size is $n$ = 30 tissues for both.

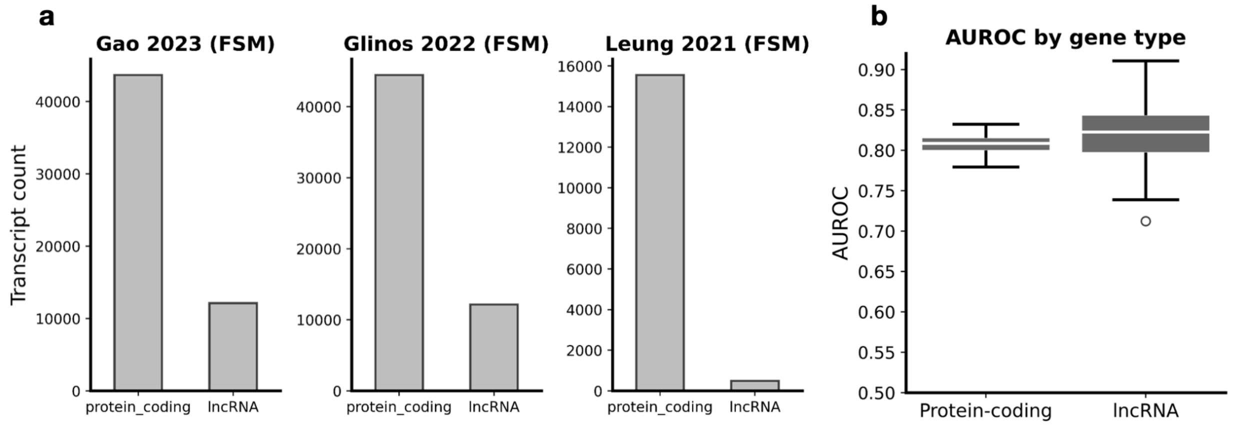


**Supplementary Fig. 5 | Otari performance by gene biotype. a**, Transcript counts (*y*-axis) for protein-coding and lncRNA genes across datasets. Bar heights indicate counts. **b**, AUROC values (*y*-axis) on the Gao et al. chromosome 8 holdout test set, stratified by gene type. Center lines indicate the median, box limits denote the 25th and 75th percentiles, whiskers extend to 1.5× the interquartile range, and open circles indicate outliers.

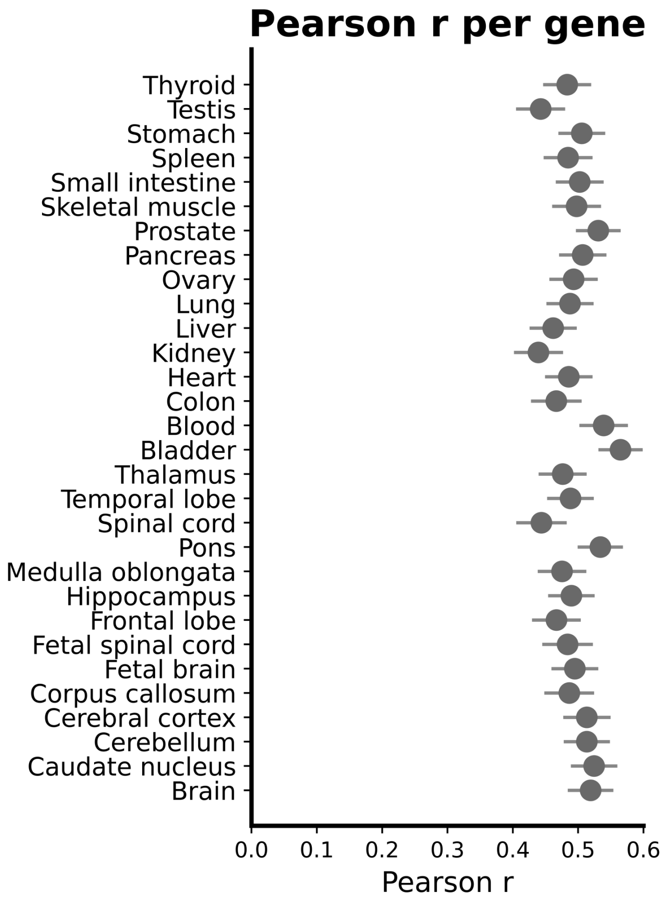


**Supplementary Fig. 6 | Per-gene prediction accuracy across tissues.** Distribution of Pearson correlation coefficients ($x$-axis) between predicted and observed isoform abundances for individual genes across 30 tissues ($y$-axis). Genes with fewer than two isoforms were excluded. Based on 2,649 transcripts in the Gao et al. chromosome 8 holdout test set. Points show means; error bars denote standard error.

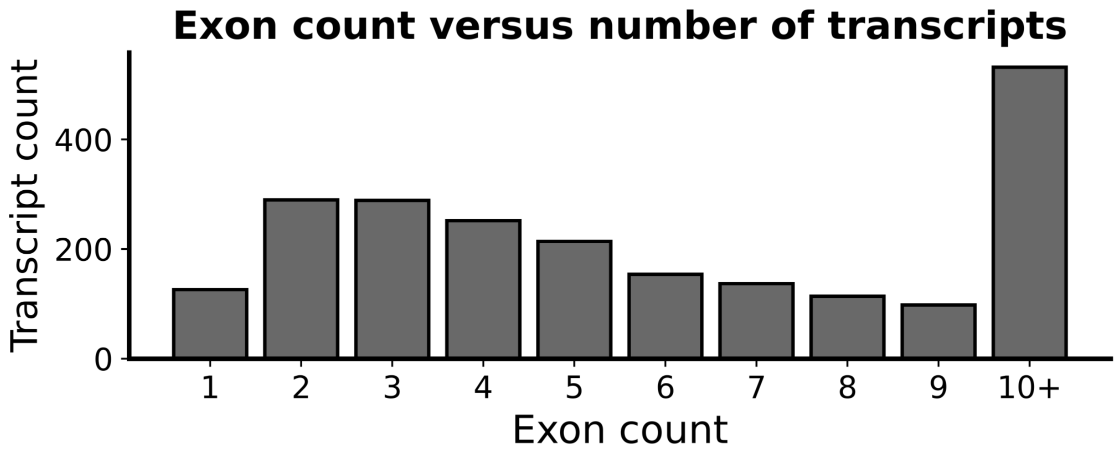


**Supplementary Fig. 7 | Transcript exon count distribution.** Transcript counts (*y*-axis) stratified by exon number (1–9 or 10+; *x*-axis) in the Gao et al. chromosome 8 holdout test set.

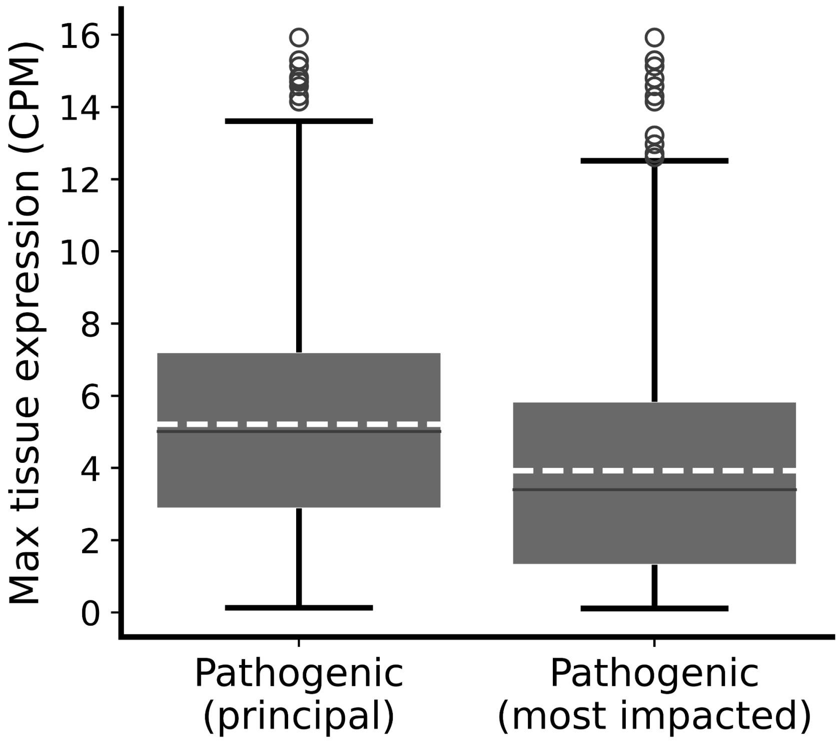


**Supplementary Fig. 8 | Expression levels of principal and top-impact transcripts.** Maximum absolute expression across tissues (*y*-axis) is shown for principal transcripts and transcripts most impacted by HGMD DFP variants (top ranked transcript per variant). Boxplot center lines indicate the median, box limits denote the 25th and 75th percentiles, whiskers extend to 1.5× the interquartile range, and open circles denote outliers. Dashed lines indicate mean expression (5.27 CPM, 3.98 CPM).

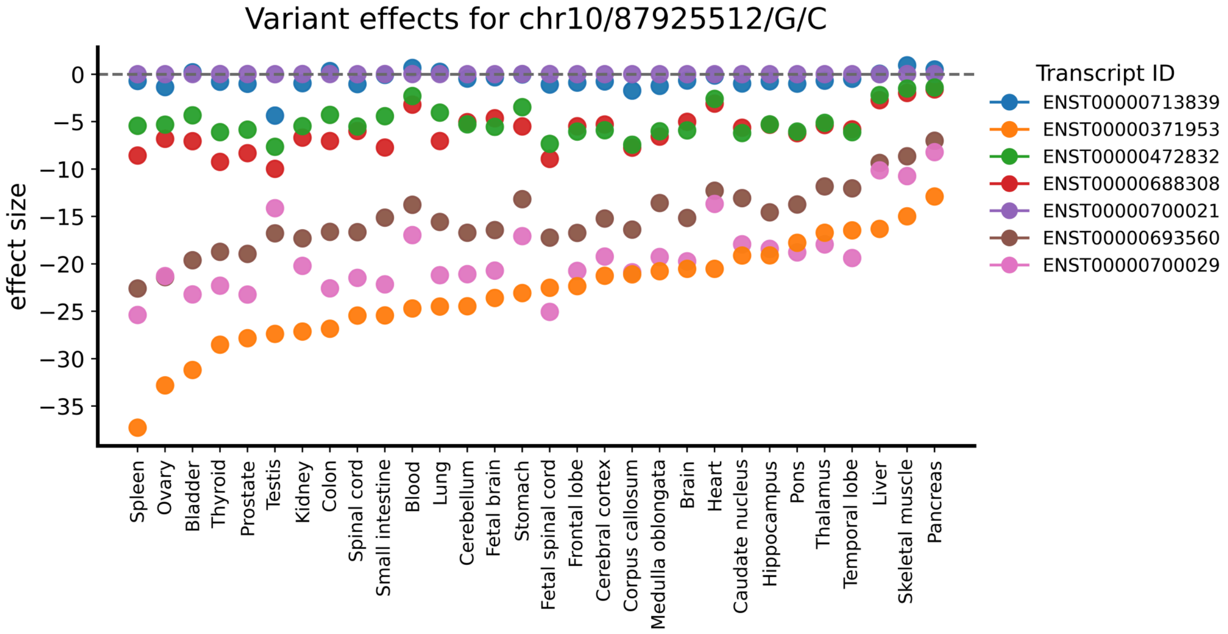


**Supplementary Fig. 9 | Predicted tissue-specific impact of rs786203847 on all GENCODE-annotated PTEN transcripts.** Predicted fold changes (*y*-axis), normalized to a background distribution (Methods), are shown across tissues (*x*-axis). Dashed line marks zero effect. Different transcripts are represented by distinct colors.

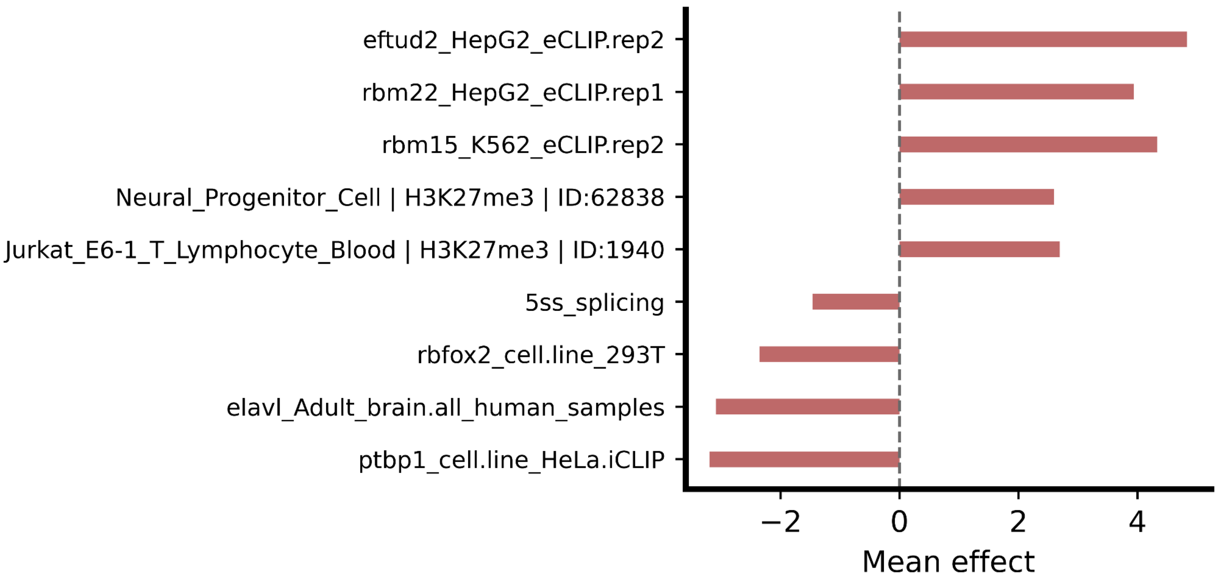


**Supplementary Fig. 10 | Regulatory features disrupted by rs786203847.** Top disrupted features were identified from the most affected nodes in PTEN transcripts. *x*-axis bar heights show mean feature effect (mean z-score computed per feature across the reference and alternative graph node attributes) and direction of effect.
